# Crystal structure and site-directed mutagenesis of circular bacteriocin plantacyclin B21AG reveals cationic and aromatic residues important for antimicrobial activity

**DOI:** 10.1101/2020.07.02.185538

**Authors:** Mian-Chee Gor, Ben Vezina, Róisín M. McMahon, Gordon J. King, Santosh Panjikar, Bernd H. A. Rehm, Jennifer L. Martin, Andrew T. Smith

## Abstract

Plantacyclin B21AG is a circular bacteriocin produced by *Lactobacillus plantarum* B21 which displays antimicrobial activity against various Gram-positive bacteria including foodborne pathogens, *Listeria monocytogenes* and *Clostridium perfringens*. It is a 58-amino acid cyclised antimicrobial peptide, with the N and C termini covalently linked together. The circular peptide backbone contributes to remarkable stability, conferring partial proteolytic resistance and structural integrity under a wide temperature and pH range. Here, we report the first crystal structure of a circular bacteriocin from food grade *Lactobacillus*. The protein was crystallised using the hanging drop vapour diffusion method and the structure solved to a resolution of 1.8 Å. Sequence alignment against 17 previously characterised circular bacteriocins revealed the presence of conserved charged and aromatic residues. Alanine substitution mutagenesis validated the importance of these residues. Minimum inhibitory concentration analysis of these Ala mutants showed Phe^8^Ala and Trp^45^Ala mutants displayed a 48- and 32-fold reduction in activity, compared to wild type. Lys^19^Ala mutant displayed the weakest activity, with a 128-fold reduction. These experiments demonstrate the importance of aromatic and cationic residues for the antimicrobial activity of plantacyclin B21AG and by extension, other circular bacteriocins sharing these evolutionarily conserved residues.

## Introduction

Bacteriocins are a group of ribosomally-synthesised antimicrobial peptides produced by both Gram-negative and Gram-positive bacteria. They generally confer antimicrobial activity against bacterial species and/or strains closely related to the bacteriocin producers. Many studies ^1-4^ have suggested that these antimicrobial peptides have potential as natural food preservative. Bacteriocins have also been identified as possible next generation antibiotics to combat multiple-drug resistant pathogens ^3,5^. In recent years, bacteriocins from Gram-positive bacteria, especially lactic acid bacteria (LAB) have attracted interest because they are generally regarded as safe (GRAS) for human consumption and are thought to have a broader antimicrobial spectrum than bacteriocins produced by Gram-negative bacteria ^6^. Furthermore, Gram-positive bacteriocins are active against foodborne pathogens, making them important candidates for controlling food spoilage and pathogenic bacteria in food and pharmaceutical industries ^6^.

Bacteriocins produced by Gram-positive bacteria are divided into three main classes: class I - modified, class II - unmodified and class III - large, heat labile ^7^. Of these, circular bacteriocins (class I) have gained attention due to their unique characteristics of high thermal and pH stability, as well as resistance to degradation by many proteolytic enzymes. The covalently linked N- and C-termini form a structurally distinct cyclic peptide backbone ^8^. There are 20 circular bacteriocins reported in the literature to date, all are between 58 to 70 amino acids in length, corresponding to approximately 5.6 to 7.2 kilodalton (kDa) in mass ^9,10^. They are clustered into two families based on sequence similarity and biochemical characteristics. Family I bacteriocins are more cationic and have higher isoelectric point (pI ∼ 10) whereas members of family II are more hydrophobic and have lower isoelectric point (pI ∼5) ^8,11^. Family I includes aureocyclicin 4185 ^12^, enterocin NKR-5-3B ^13^, amylocyclicin ^14^, amylocyclicin CMW1 ^15^, enterocin AS-48 ^16^, bacA ^17^, carnocyclin A ^18^, circularin A ^19^, thermocin 458 ^20^, garvicin ML ^21^, lactocyclicin Q ^22^, leucocyclicin Q ^23^, pumilarin ^9^, uberolysin ^24^ and cerecyclin ^25^. These bacteriocins are found across a wide range of Gram-positive genera. Family II are generally found in *Lactobacillus, Staphylococcus* and *Streptococcus* species, and includes gassericin A/reutericin 6 ^26^, butyrivibriocin AR10 ^27^, acidocin B ^28^, paracyclicin ^29^, plantaricyclin A ^30^ and plantacyclin B21AG ^31-33^.

Until now, the only bacteriocin crystal structure reported was that of enterocin AS-48, which was solved by single isomorphous replacement with anomalous scattering (SIRAS) ^34^. By comparison there are many Nuclear Magnetic Resonance (NMR) structures of bacteriocins ^35^. For example, NMR solution structures have been reported for family I enterocin AS-48, carnocyclin A, enterocin NKR 5-3B and family II, acidocin B. Despite low sequence similarity, all the NMR and crystal structures demonstrate a conserved structural motif consisting of four to five α-helices encompassing a hydrophobic core, with the C-teminus and N-terminus ligation occurring within an α-helix secondary structure ^11^. The circular nature of these proteins contributes to remarkable stability against physical stresses ^8,36^. The conformational and thermal stability of the circular enterocin AS-48 compared to its linear counterpart, AS_10/11_ obtained by limited proteolysis have been demonstrated previously ^37^. For example, the linear AS_10/11_ has 35 % lower α-helical content compared with the native circular protein (measured by Far-UV circular dichroism (CD)). Linear AS_10/11_ is less compact and rigid compared to circular AS-48 ^38^. Linear AS_10/11_ at pH 2.5 showed a low cooperativity of thermal unfolding and reduced stability compared to the circular AS-48, which was shown to unfold at 102 °C ^39,40^. The linear AS_10/11_ retained some antimicrobial activity although it was 300 times lower than the circular AS-48, suggesting that circularisation is not essential for bactericidal activity but more importantly for stabilisation of the three-dimensional structure of the bacteriocin ^37^. Similarly, the linear version of AS-48, Trp70Ala mutant has just 18 % helical structure compared to 72 % in the circular AS-48, as measured by far-UV CD. Circular AS-48 is extremely stable in comparison with its linear counterpart, with no sigmoidal transition observed between 25 and 95 °C, whereas linear Trp70Ala mutant had a melting temperature of 61 °C ^41^. These structurally stable circular bacteriocins and their mechanism of circularisation are of interest because of their biotechnological properties and applications. Proteins with new or improved features including enhanced stability could be generated through protein engineering, though this requires a deep understanding of bacteriocin structure-function relationships ^18,38^.

We previously identified a circular bacteriocin plantacyclin B21AG from *Lactobacillus plantarum* B21. This family II circular bacteriocin has antimicrobial activity against foodborne pathogens including *Listeria monocytogenes* and *Clostridium perfringens* and other closely related *Lactobacillus* species ^31^. We have also cloned the plantacyclin B21AG gene cluster into a plantacyclin B21AG-negative strain and used this system for its expression ^42^. In the present study, we report the crystal structure of plantacyclin B21AG. Using this crystal structure, we identified by structural bioinformatics several conserved residues and showed via site-directed mutagenesis that these residues contribute to bactericidal function. Only one crystal structure has been reported for an enterocin AS-48, from bacteriocin family I which is evolutionarily quite distinct from family II circular bacteriocins ^10^. Elucidation of the structure of plantacyclin B21AG could therefore provide new information in understanding the structure and function of these fascinating proteins. To the best of our knowledge, this is the first report of a crystal structure of a family II circular bacteriocin, and the first of a bacteriocin produced by a food grade LAB.

## Results and Discussion

### Crystal structure of plantacyclin B21AG

Crystals of plantacyclin B21AG suitable for structure determination grew from hanging drop vapour diffusion using a well solution of 1.1 M sodium malonate, 0.1 M HEPES buffer pH 7.0 and 0.5% v/v Jeffamine® ED-2003. A crystal from these conditions diffracted to 1.8 Å resolution and belong to the space group C222_1_. Crystallographic statistics for data collection/processing and refinement are shown in Table 1. The crystal structure diffraction data were phased by molecular replacement using a theoretical α-helix (8 residues long polyalanine) as a search model. The structure was refined to a final R_work_/R_free_ of 0.169/0.215 at 1.8 Å resolution. There are two independent molecules of plantacyclin B21AG in the asymmetric unit. The structures of these two molecules (Chain A and Chain B) are similar to each other, with a C^α^ backbone root-mean-square deviation, RMSD of 0.56 Å across 58 C^α^ atoms (Figure 1a).

**Table 1.** Data collection and refinement statistics for the crystal structure of plantacyclin B21AG.

|  |  |
| --- | --- |
| **Data collection** |  |
| Space group | C222<sub>1</sub> |
| Cell dimensions |  |
| a, b, c (Å) | 44.13, 93.17, 49.49 |
| $\alpha$, $\beta$, $\gamma$ (°) | 90, 90, 90 |
| Resolution (Å) | 46.58 – 1.80 (1.84 – 1.80)<sup>a</sup> |
| No. observations | 128402 |
| No. reflections | 9770 |
| R<sub>merge</sub> | 0.074 (0.730) |
| R<sub>pim</sub> | 0.030 (0.306) |
| *I* / $\sigma I$ | 15.5 (2.4) |
| Completeness (%) | 99.8 (96.7) |
| Redundancy | 13.1 (12.3) |
| **Refinement** |  |
| Resolution (Å) | 46.58 – 1.80 |
| No. reflections | 9770 |
| *R*<sub>work</sub> / *R*<sub>free</sub> | 16.9% / 21.5% |
| No. atoms |  |
| Protein | 802 |
| Ligand/ion | 7 |
| Water | 56 |
| *B*-factor |  |
| Protein | 26.1 |
| Ligand/ion | 21.6 |
| Water | 39.2 |
| R.m.s. deviations |  |
| Bond lengths (Å) | 0.019 |
| Bond angles (°) | 1.697 |

The number of crystals used for structure determination and refinement is one.
<sup>a</sup> Highest resolution shell is shown in parenthesis.

**Figure 1.**
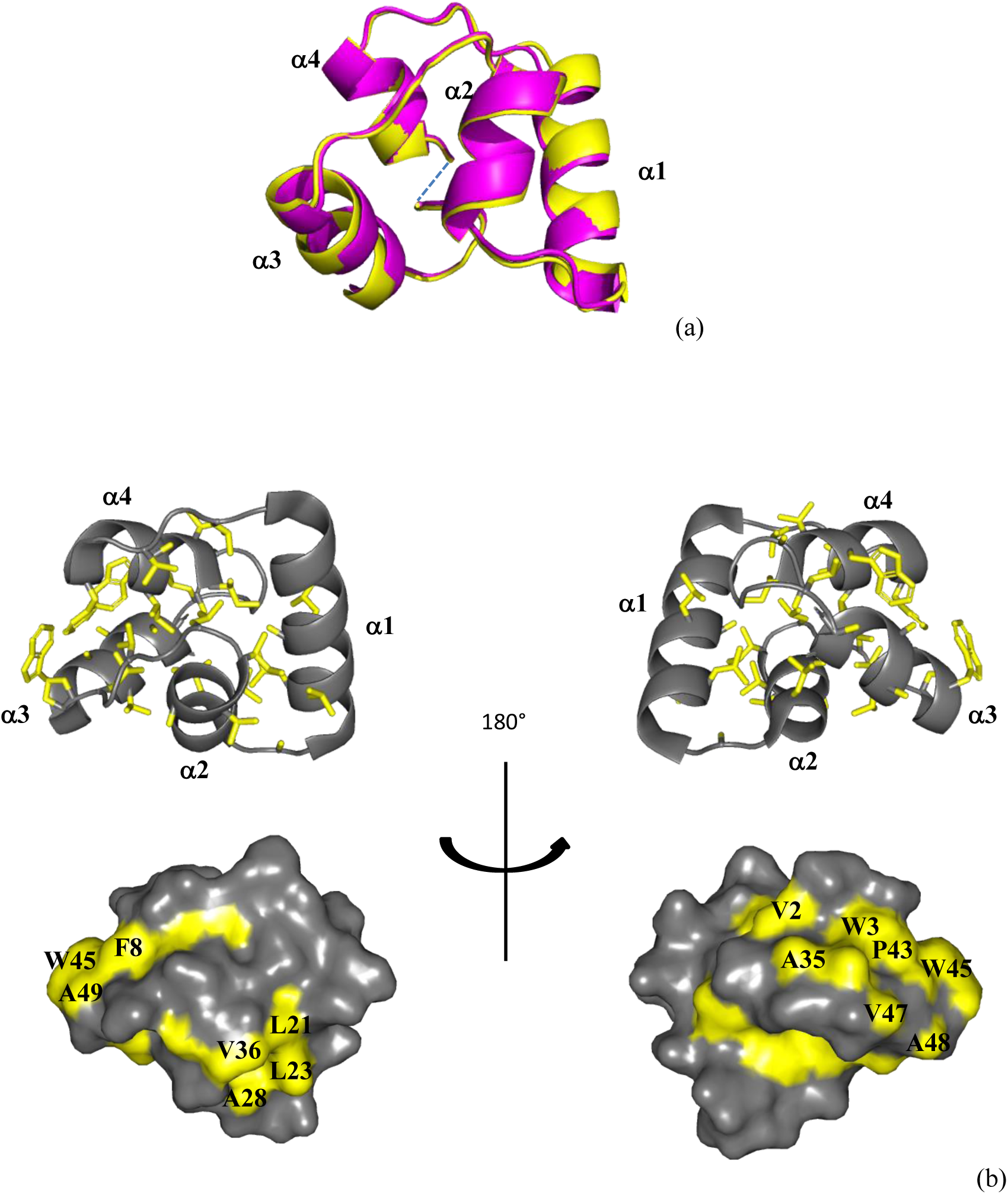
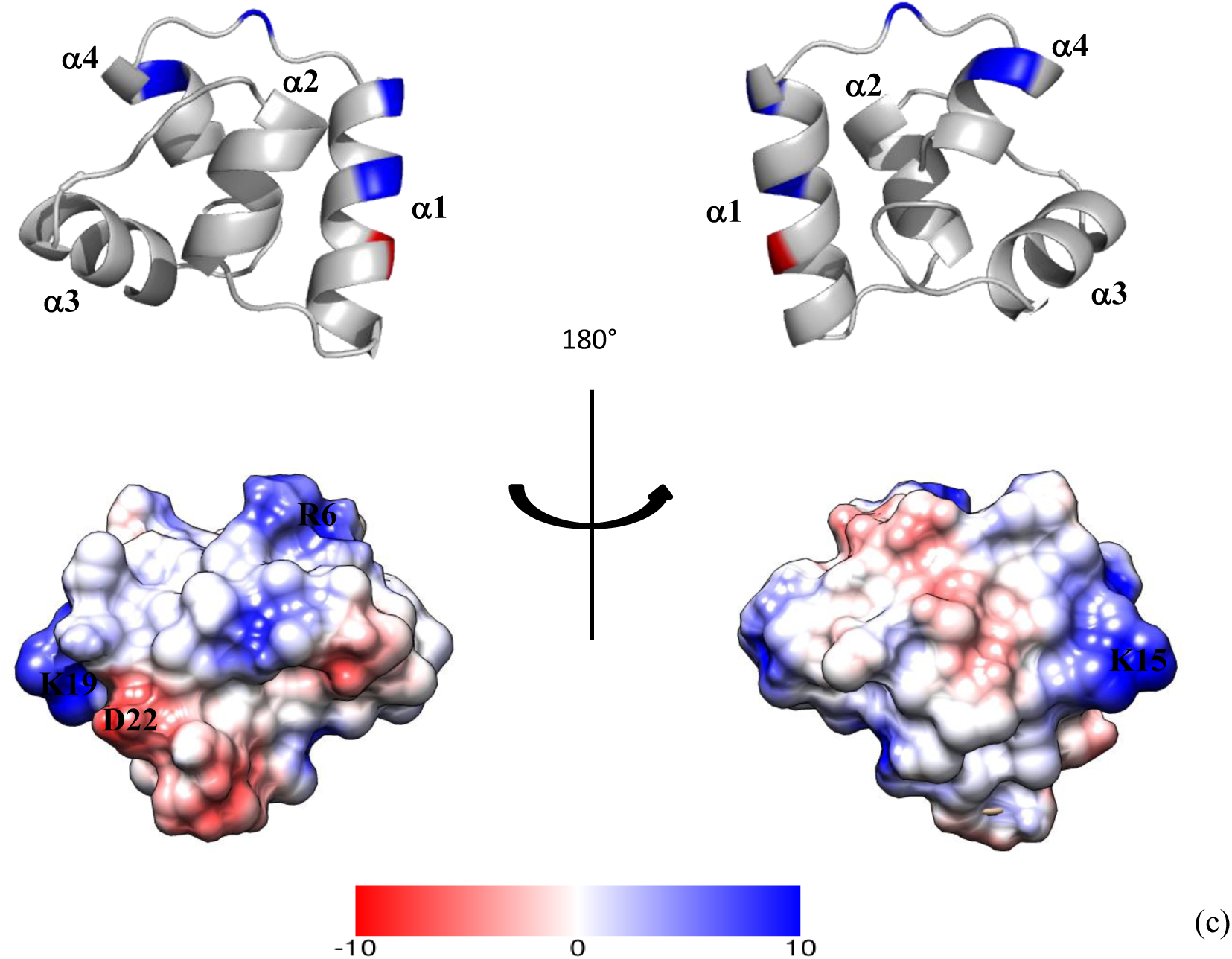
Structural characteristics of plantacyclin B21AG. (a) Superposition of Chain A (magenta) and Chain B (yellow) of the crystal structure of plantacyclin B21AG. The blue dashed line indicates the N-to-C cyclisation point. (b) Hydrophobicity of plantacyclin B21AG. In this figure, the side chain atoms of hydrophobic residues (Ala, Ile, Leu, Phe, Pro, Gly, Trp and Val) are coloured in yellow. All other atoms of the same residues and all atoms of non-hydrophobic residues are shown in grey. Upper panel: Ribbon diagram illustrating the compact, hydrophobic core of plantacyclin B21AG. Hydrophobic side chains are drawn as sticks; Lower panel: Surface representation showing the solvent exposed hydrophobic side chains. (c) Ribbon diagram (upper panel) and electrostatic potential (lower panel) depicting amphipathicity of plantacyclin B21AG. Electrostatic potential calculated by APBS function^82^ using PDB2PQR version 2.1.1 ^83^ and surface map generated with Chimera 1.14 ^84^. Cationic, anionic and neutral residues are depicted in blue, red and white, respectively. Key residues are labelled.

The structure of plantacyclin B21AG is characterised by four α-helices, i.e. Thr^14^ – Ser^26^ (α1), Leu^30^ – Leu^38^ (α2), Gly^44^ – Ala^52^ (α3) and Gly^54^ – Phe^8^ (α4). Helix α1 and helix α4 are linked covalently between Ile^1^ and Ala^58^ in the middle of helix α4, creating a circular backbone (Figure 1a). The four α-helices in the crystal structure are connected by loops of 1 to 6 residues and enclose a hydrophobic core in a tightly folded globular arrangement (Figure 1b). Together, the loops contain five surface exposed glycine residues i.e. Gly^9^, Gly^27^, Gly^39^, Gly^54^ and Gly^55^. Glycines are often found in flexible loop regions as they do not contain a side chain, increasing the conformational flexibility (demonstrated by NMR solution structures of other circular bacteriocins) ^13,28,43^. The hydrophobic core residues are Ile^1^, Ile^4^, Ala^5^, Val^10^, Leu^12^, Ala^20^, Leu^24^, Leu^30^, Val^33^, Ala^34^, Ile^37^, Leu^38^, Val^40^, Leu^42^, Ala^46^, Ala^50^ and Ala^58^. All of these residues have <20% solvent exposure calculated based on GETAREA^44^. The second, third and fourth α-helices are predominantly hydrophobic, with the linked amino acids at the N- and C-termini in helix α4 buried in the core of the structure. In contrast, helix α1 is amphipathic, with hydrophilic residues (Thr^14^, Gln^18^, Ser^25^ and Ser^26^) exposed at the surface of the protein.

The protein has an overall net charge of +3, with four cationic residues located in helix α1 (Lys^15^ and Lys^19^), helix α4 (Arg^6^) and the loop connecting α1/α4 (His^11^), and one anionic residue located in helix α1 (Asp^22^) (Figure 1c, upper panel). These charged residues are all displayed on the surface of the plantacyclin B21AG structure. The electrostatic potential surface map of plantacyclin B21AG reveals two cationic patches separated by an anionic strip (Figure 1c, lower panel).

### Structural comparisons of circular bacteriocins

To better understand the similarities and differences between known structures of circular bacteriocins, structural comparisons were performed between plantacyclin B21AG and other bacteriocin 3D structures elucidated to date, namely enterocin AS-48, carnocyclin A and enterocin NKR-5-3B (family I) and acidocin B (family II). We note that the acidocin B structure was determined using the bacteriocin embedded in SDS micelles ^28^, whereas the other bacteriocin structures were determined under aqueous conditions ^13,34,43^. These circular bacteriocins adopt a common 3D structure comprising four or five helices folded into a globular bundle enclosing a hydrophobic core ^11^ (Figure 2, left panel) ^13,28,34,43^. The orientation of the helices is similar within the members of each family, with the exception of enterocin AS-48 which has five rather than four α-helices. Notably, the circularisation point for all these bacteriocins is located within a helical structure which contains mostly hydrophobic residues ^36^. Analysis of solvent exposed residues using GETAREA^44^ reveals that the N- to C-linkage of all these bacteriocins except the micelle embedded acidocin B is buried in the protein core (the ratio of side chain surface area to the average solvent-accessible surface area of the first and last amino acid, respectively is less than 20%). Unlike saposins and saposin-like peptides which are stabilised by disulfide bonds between cysteine residues ^45^, the bacteriocin helical fold appears to be stabilised by hydrophobic side chain interactions upon peptide circularisation ^43^. Martin-Visscher, et al. ^43^ proposed that hydrophobic residues close to the N-C linkage may play a role in the interaction between the linear peptide and the cyclisation enzyme, helping to bring the termini into close proximity for cyclisation to take place. A combination of circular backbone and hydrophobic core is thought to contribute to the stability of circular bacteriocins ^13^.

**Figure 2.**
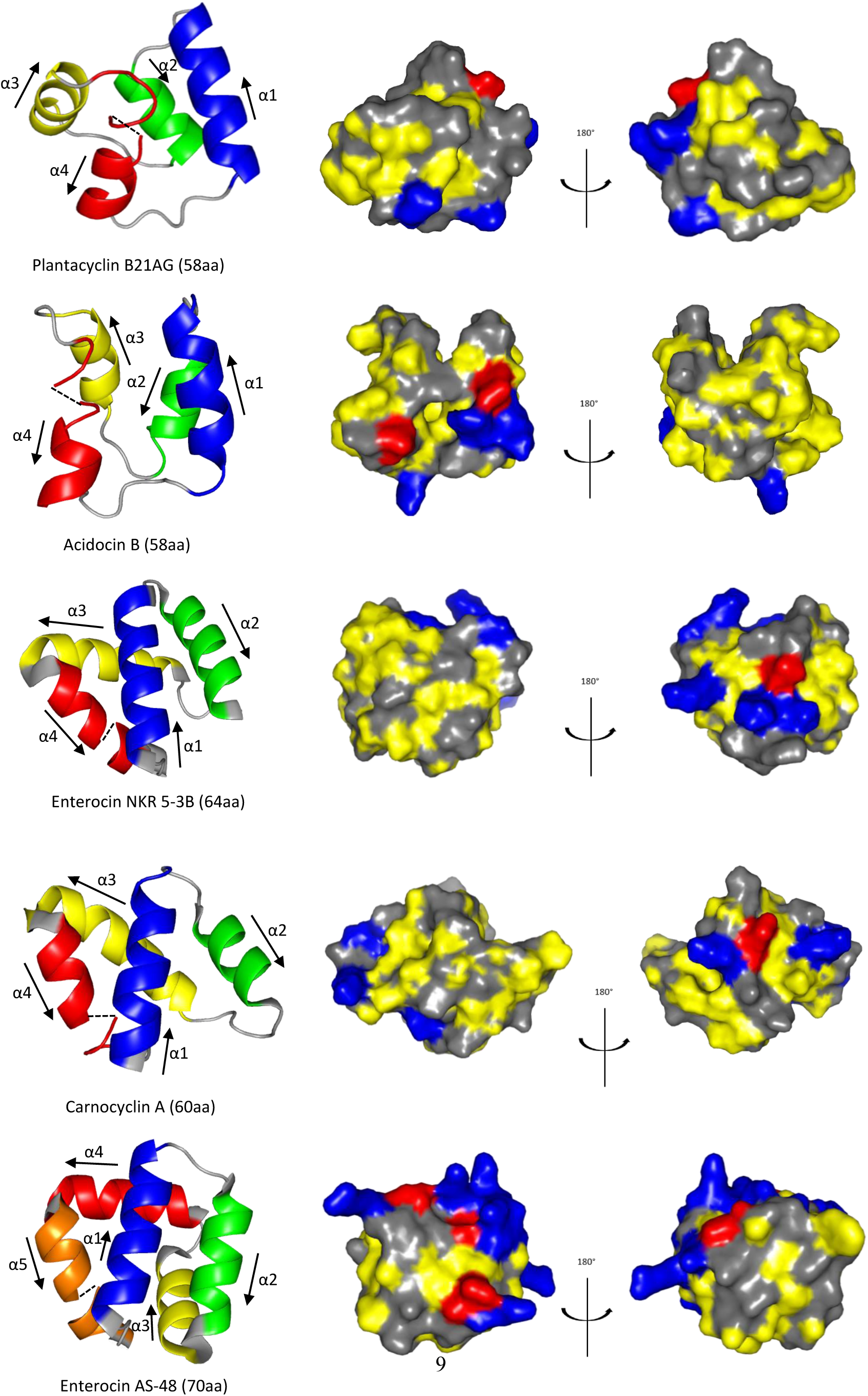
Three dimensional structural features of circular bacteriocins. Left panel: The helical orientation of the four/five α-helices is shown as α1 (blue), α2 (green), α3 (yellow), α4 (red) and α5 (orange). The black arrows indicate the orientation of the α helices. The dotted black lines represent the linkage between the N- and C-terminal. Right panel: Surface representation of the protein structures depicting side chain atoms of hydrophobic residues (yellow), cationic residues (blue), anionic residues (red) and polar residues (grey).

### Surface characteristics of circular bacteriocins

The molecular surface of the circular bacteriocins is amphipathic: one face of the molecule has cationic and anionic patches, and the other surface is uncharged or hydrophobic (Figure 2, right panel). Generally, family I circular bacteriocins are more cationic than family II circular bacteriocins (Table 2). Noteworthy is the cationic region located close to the N- and C-ligation point (α1 and α4). This cationic region is present in enterocin AS-48, carnocyclin A and enterocin NKR-5-3B. Eight lysine residues are located along the stretch of α4 and α5 in enterocin AS-48. Similarly, four lysine residues are located along α3 and α4 of carnocyclin A and enterocin NKR-5-3B. The data presented in Figure 2 and Table 2 suggest that the charge distribution is conserved among circular bacteriocins ^13^. The cationic surface patches are thought to be involved in an initial electrostatic interaction between the peptide and the negatively charged phospholipid bilayer of target cell membranes ^8,13,36^. Binding of the cationic regions onto the target membrane may result in destabilisation of the membrane, potentially enabling peptide insertion ^43^. However, it has also been demonstrated that electrostatic interaction alone is insufficient for antimicrobial activity. A 21-residue peptide fragment of enterocin AS-48 containing the cationic putative membrane interacting region exhibited competitive membrane binding ^46^ but did not show antibacterial activity, suggesting that other physicochemical properties of the bacteriocins may be required for antimicrobial action.

**Table 2.** Physical properties of the circular bacteriocins with solved structures.

| Bacteriocin | Leader peptide (aa) | Mature peptide (aa) | MW (Da) | pI<sup>a</sup> | Net charge | Hydrophobicity<sup>a</sup> (GRAVY Index)<sup>b</sup> | Producer organism | Reference |
| --- | --- | --- | --- | --- | --- | --- | --- | --- |
| **Family I** |  |  |  |  |  |  |  |  |
| Enterocin AS-48 | 35 | 70 | 7150 | 10.1 | +6 | 0.539 | *Enterococcus faecalis* AS-48 | Samyn, et al. <sup>16</sup> |
| Carnocyclin A | 4 | 60 | 5862 | 10.0 | +4 | 1.058 | *Carnobacterium maltaromaticum* UAL 307 | Martin-Visscher, et al. <sup>18</sup> |
| Enterocin NKR-5-3B | 23 | 64 | 6317 | 9.9 | +5 | 0.953 | *Enterococcus faecium* NKR-5-3 | Himeno, et al. <sup>13</sup> |
| **Family II** |  |  |  |  |  |  |  |  |
| Plantacyclin B21AG | 33 | 58 | 5668 | 10.0 | +3 | 1.002 | *Lactobacillus plantarum* B21 | Golneshin, et al. <sup>31</sup> |
| Acidocin B | 33 | 58 | 5622 | 6.8 | +1 | 1.036 | *Lactobacillus acidophilus* M46 | Acedo, et al. <sup>28</sup> |

<sup>a</sup>Predicted for the linear bacteriocin using ExPASy ProtParam Tool (https://web.expasy.org/cgi-bin/compute_pi/pi_tool). <sup>b</sup>Grand Average of Hydrophathy.

Most of the bacteriocin anionic residues are located in close proximity to the cationic patches (Figure 2, right panel). Whilst the role of anionic residues are generally overlooked in the context of circular bacteriocins, studies have shown that mutation of anionic residues in other classes of bacteriocins and/or antimicrobial peptides resulted in reduced bactericidal potency and altered target cell specificity ^47,48^. Replacing aspartic acid at position 17 with the glutamic acid strongly enhanced the antimicrobial activity of the pediocin-like bacteriocin sakacin P. This result suggests that the anionic residues interact in a structurally specific and restricted manner with a cationic region on the target cell or on the peptide itself ^49^. Given that bacteriocins generally kill through disruption of proton motive force ^50^ or through the Grotthuss mechanism of gramicidin ^51^, placement of negative charges near the pore may facilitate pore selectivity and cation flow out of the target cell by reducing the electrostatic energy profile and increasing selectivity for divalent cations such as Ca^2+^ ^52^. These findings suggest that anionic residues are as important as cationic residues in terms of target specificity of the bacteriocins.

The Grand Average of Hydropathy (GRAVY) Index ^53^ calculation reveals that family II circular bacteriocins are more hydrophobic than those in family I - except Circularin A (Table 2) The hydrophobicity of plantacyclin B21AG is evident in that it selectively dissolves into butanol fractions during purification. Some hydrophobic residues are exposed at the surface of the molecules (Figure 2, right panel). The hydrophobicity of these peptides is thought to be crucial for creating pores in the bacterial membrane, especially for family II bacteriocins as they are less cationic ^28,45^ (Table 2). Permeation of the cell membrane by these peptides causes leakage of ions, dissipation of membrane potential and eventually cell death ^8^. The molecular mode of action of enterocin AS-48 has been widely studied and is the best understood of the circular bacteriocins. A model has been proposed ^34^ such that the peptide in the form of a water-soluble dimer approaches the membrane surface of target bacteria through electrostatic interaction. Upon membrane interaction, each protomer within the water-soluble dimer rotates 90° and rearranges such that the hydrophobic helices become solvent accessible. The transition of AS-48 from water-soluble dimer to membrane-bound dimer allows the bacteriocin to insert into the bacterial membrane. This insertion alters the membrane potential, causing pore formation and cell leakage ^34,46,54^. Collectively, these studies ^34,46,54^ have shown that cationic patches and hydrophobic patches play an important role in the mechanism of action of bacteriocins.

### Structural alignment of circular bacteriocins

Structural alignment using the align function in PYMOL ^55^ revealed that the three-dimensional structure of family II plantacyclin B21AG aligned well with family I enterocin NKR5-3B, with a r.m.s.d of 2.47 Å across 30 C^α^ atoms. For both structures, the helices α1, α2 and α4 are similar in length and align in parallel in a similar orientation. α3 of enterocin NKR5-3B is twice the length of α3 of plantacyclin B21AG but is oriented similarly (Figure 3). The alignment of plantacyclin B21AG with enterocin NKR5-3B suggests that family I and family II bacteriocins could share a conserved three-dimensional structure with a very similar core fold, despite low sequence identity (15.6%) (Figure 4). In contrast, there is poor structural alignment between plantacyclin B21AG and enterocin AS-48 with r.m.s.d of 4.54 Å across 45 C^α^ atoms (Figure 3) (sequence identity 11.4%, Figure 4). AS-48 also has a longer sequence than plantacyclin B21AG (70 residues compared with 58) and forms five α-helices rather than four (Figure 2). Similarly, plantacyclin B21AG and carnocyclin A do not align well, with r.m.s.d of 4.33 Å across 45 C^α^ atoms (Figure 3) (sequence identity 16.7%, Figure 4).

**Figure 3.**
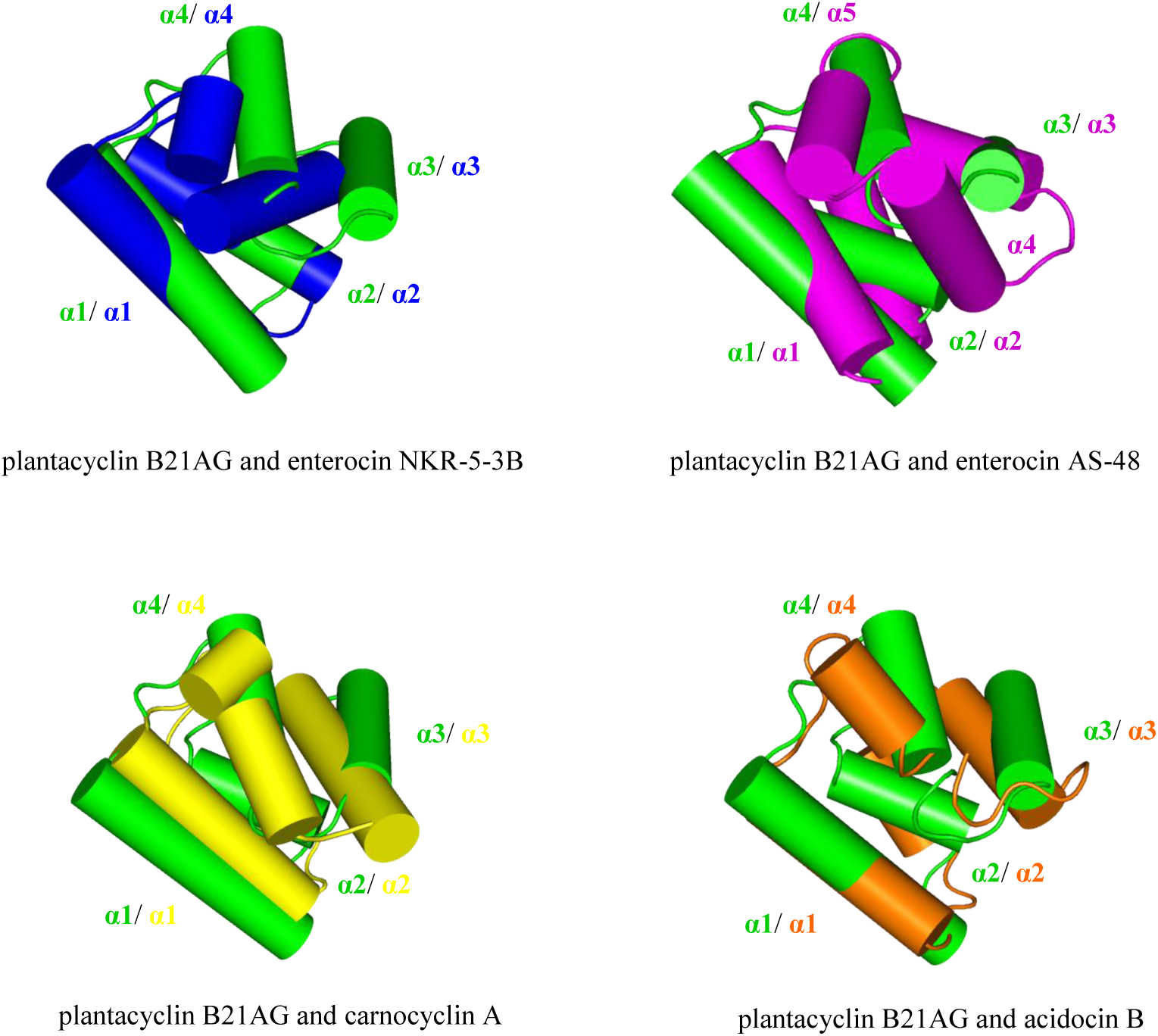
Structural alignment of plantacyclin B21AG (green) with enterocin NKR-5-3B (blue), enterocin AS-48 (magenta), carnocyclin A (yellow) and acidocin B (orange). The helices are labelled accordingly using plantacyclin B21AG as reference.

**Figure 4.**
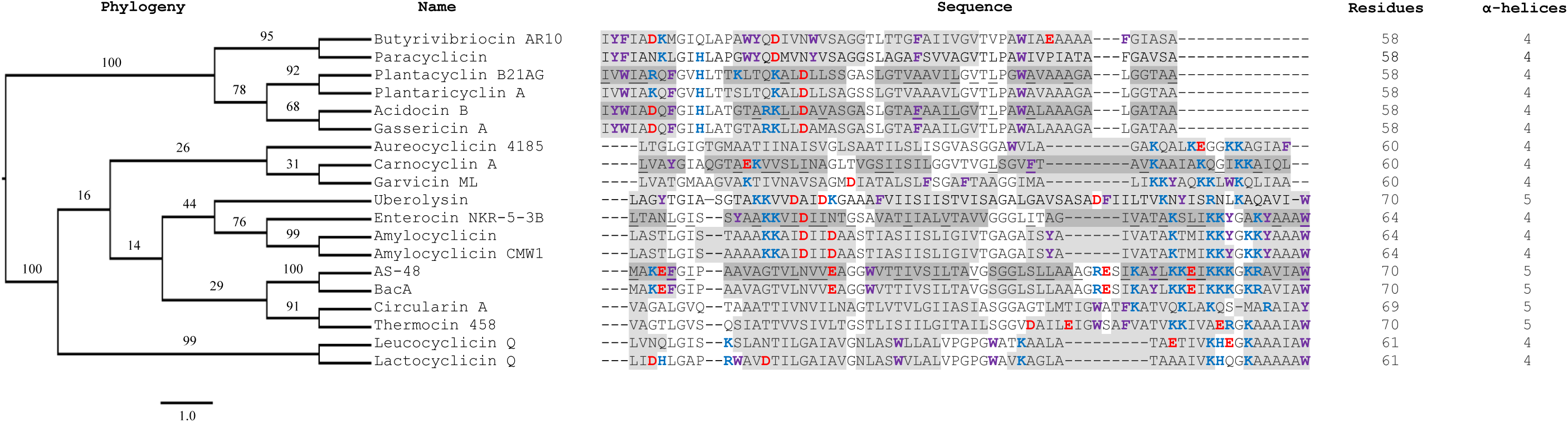
Phylogenetic tree showing the evolutionary relationship between the mature circular bacteriocin sequences, and their aligned sequences. Blue shows cationic residues. Red shows anionic residues. Purple shows aromatic residues. Dark grey highlighting shows α-helices confirmed via structural information. Light grey highlighting shows α-helices predicted by sequence alignment and secondary structure prediction. Underlining shows residues buried in protein core in experimentally determined structures. A. Sequence length of each protein is shown in “residues” column and number of known or predicted α-helices is shown in column labelled α-helices.

Plantacyclin B21AG shares 65% sequence identity with acidocin B (Figure 4), and both belong to the circular bacteriocin family II. As indicated above, the NMR structure of acidocin B was determined in the presence of SDS micelles ^28^. Not surprisingly, the structure of plantacyclin B21AG (determined in aqueous conditions) did not align well with that of acidocin B, giving a r.m.s.d. of 4.95 Å across 57 C^α^ atoms (Figure 3). Compared with aqueous conditions, SDS micelles may better mimic the bacterial membrane. The comparison of the NMR solution structure of acidocin B and the crystal structure of plantacyclin B21AG may therefore provide clues to the structural changes that occur in bacteriocin family II proteins upon membrane interaction.

### Sequence analysis of circular bacteriocins

To extend the structural comparison, the sequence of plantacyclin B21AG was aligned against the sequences of 17 bacteriocins that have been previously characterised as circular. In each case, circularisation had been confirmed structurally or validated via independent experiments including electrospray ionization time-of-flight mass spectrometry (ESI-TOF MS) and peptide sequencing. Amino acid sequence alignment of these circular bacteriocins show two families within the class I circular bacteriocins (Figure 4), which is consistent with recent phylogenetic analysis ^10^. Secondary structure prediction suggests that these circular bacteriocins share considerable structural similarity despite sequence variation both within and between the two families. The results suggest that a sequence length of 69 – 70 residues gives a five α-helical structure whereas a bacteriocin sequence with fewer than 69 residues gives a four helical structure. Jpred secondary structure prediction ^86^ matched the experimentally determined secondary structure despite the large sequence diversity of the circular bacteriocins.

The position in the sequence of key amino acids such as charged, buried and aromatic residues appears to be conserved across the circular bacteriocin, which may hint at a key role for these residues in the protein antimicrobial activity. Specifically, the cationic residues located in helix α3 and α4 (helix α4 and α5 in the case of enterocin AS-48) in family I are predominantly but not exclusively lysine. In family II, histidine at position 11 and lysine at position 19 are highly conserved. These residues generally appear in polycationic clusters within the first, second, fourth/fifth α-helices. Upon N- to C-termini ligation, these cationic residues are brought into close proximity.

Anionic residues in the circular bacteriocins are generally conserved, with almost all being found in close proximity to the membrane-interacting polycationic region and aromatic residues (Figure 4). This matches the pattern observed with the structurally determined circular bacteriocins (Figure 2, right panel). The N-C linkage point is buried in the hydrophobic core in the majority of experimentally determined structures of circular bacteriocins. In family I, the N- and C-terminal residues are generally conserved, sharing common characteristics. For example, the N terminus of family I utilises most commonly a leucine, then valine and methionine. These are all hydrophobic residues which are similar in size (117 – 149 Da). The C terminus consists of an aromatic residue, either a tryptophan, tyrosine or phenylalanine. Carnocyclin A and garvicin ML are exceptions to this pattern, having a leucine and alanine, respectively at the C-terminus. Instead, a conserved aromatic residue is present in the post-ligation site for carnocyclin A and in the pre-ligation site in garvicin ML. The termini of family II are also highly conserved in both sequence and residue characteristics. The N-terminal residue is often isoleucine, or an aromatic residue, or a valine while the C terminus is an alanine in every case.

Aromatic residues are also highly conserved in family I and II and in addition to the C terminus of almost every family I sequence are found near the termini of predicted or confirmed α-helices. Tryptophan, phenylalanine and tyrosine that flank transmembrane-associated helices in other proteins are thought to facilitate membrane penetration ^56,57^. This may also be the case for the circular bacteriocin sequences (Figure 4). The presence of conserved aromatic residues near α-helices in circular bacteriocins may suggest they assist in cell membrane permeation and pore formation. For example, Trp^24^ is essential for the biological activity of AS-48 and is located in a hydrophobic region that interacts with the membrane ^58^. Tryptophan has a preference for the interfacial region of lipid bilayers and allows penetration of them ^59^. Trp^24^ is essential for antimicrobial killing activity of gramicidin ^60^, specifically for channel formation and conductance ^61^ and altering energy profiles for ion permeation through long-range electrostatic interactions ^57^. Tyrosine and phenylalanine can also have a similar effect ^62^, with phenylalanine increasing membrane permeability of proteins ^63^.

### Mutagenesis of key residues in plantacyclin B21AG

To confirm the importance of specific residues, we performed site-directed mutagenesis on aromatic and cationic residues within the mature sequence of plantacyclin B21AG. These were non-synonymously substituted with alanine (Ala) to test their importance for antimicrobial activity. Figure 5 shows the location of the three altered residues, Phe^8^, Lys^19^ and Trp^45^. Other Ala substitution mutants (Arg^6^ and His^11^) expression plasmids were constructed, however despite multiple attempts to transform the DNA into *L. plantarum* WCFS1, no transformants were recovered. This could be partly due to bacteriocin toxicity to recombinant host. In general, transformation into *L. plantarum* WCFS1 was inefficient, requiring many attempts for each construct.

**Figure 5.**
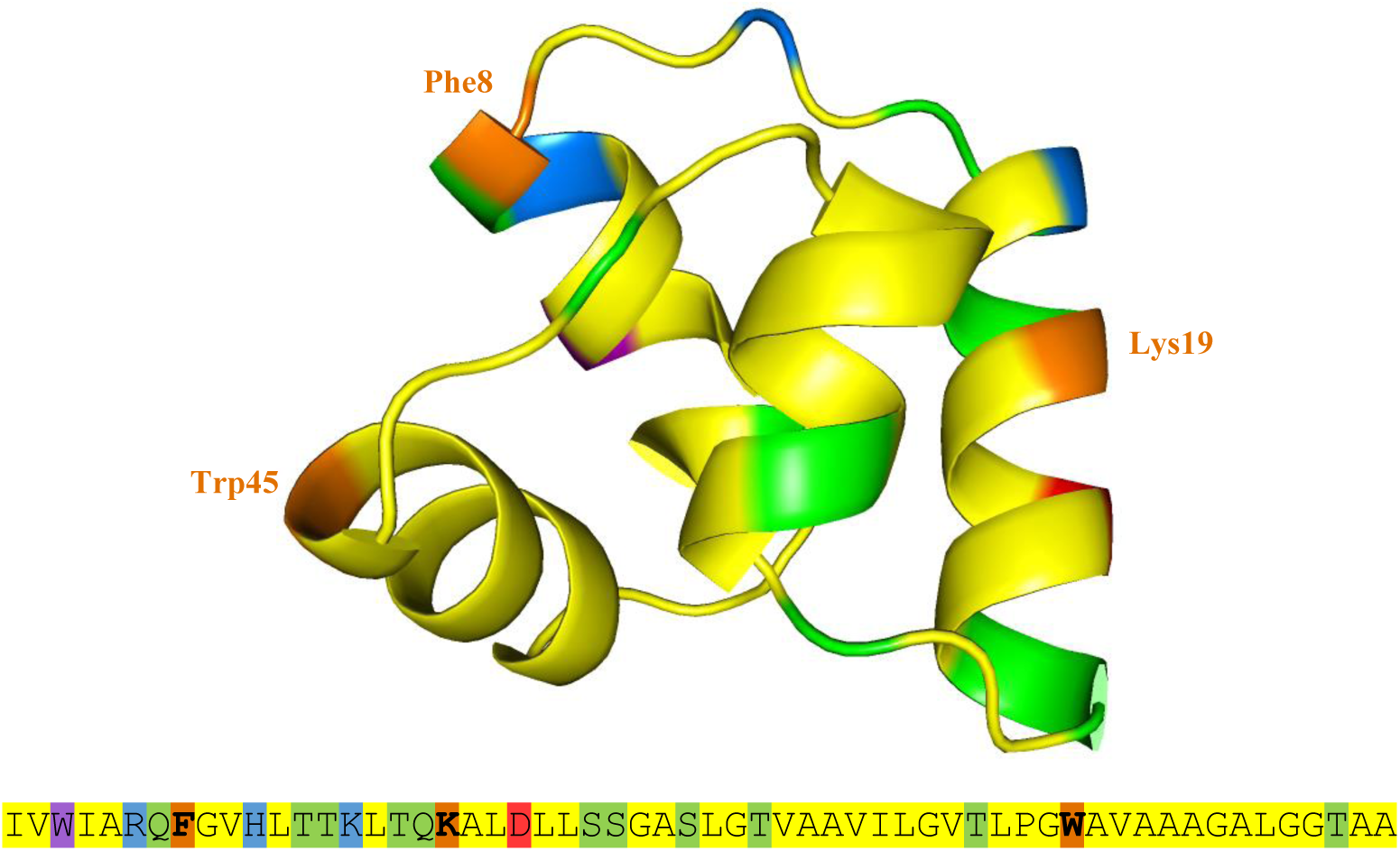
3D crystal structure of plantacyclin B21AG mutagenesis location sites visualised in PyMOL with the corresponding amino acid sequence below. Orange shows the three residues (Phe^8^, Lys^19^ and Trp^45^) non-synonymously substituted with Ala. Colour indicates the positions of hydrophobic (yellow), polar (green), aromatic (purple), cationic (blue) and anionic (red) residues.

The MIC assay results (Table 3) showed plantacyclin B21AG-WT had a value of at least 5.2 ng/μL against indicator strain *L. plantarum* A6, indicating potent antimicrobial activity against a strain closely related to the bacteriocin producer. By contrast the MIC values of each variant increased, indicating that the antimicrobial activity is reduced. The aromatic residues Phe^8^ and Trp^45^ appeared to be important for MIC activity; when replaced with Ala mutants the MIC activity was reduced by 48 and 32-fold, respectively. Mutation of aromatic residues in AS-48 also increased MIC across a range of indicator strains by a minimum of 28-fold for *Listeria monocytogenes* 4032 to 168-fold for *Enterococcus faecalis* S-47 ^58^, supporting the notion of a similar role for these residues, despite distinct phylogenetic origins (Figure 4).

**Table 3.** Results of MIC antimicrobial assay against indicator strain *L. plantarum* A6.

| *L. plantarum* WCFS1 sample | MIC<sup>a</sup> (ng/μL) | Fold-reduction of antimicrobial activity |
| --- | --- | --- |
| Plantacyclin B21AG-WT | 5.2 ± 0.0 | N/A |
| Plantacyclin B21AG-Phe<sup>8</sup>Ala | 250 ± 0.0 | 48 |
| Plantacyclin B21AG -Lys<sup>19</sup>Ala | 666.7 ± 0.0 | 128 |
| Plantacyclin B21AG -Trp<sup>45</sup>Ala | 166.7 ± 0.0 | 32 |
| Untransformed (-) | N/A | N/A |
| 10 mM ammonium acetate (-) | N/A | N/A |

MIC values are mean ± SEM of two biological replicates.
<sup>a</sup>Abbreviations: MIC, minimum inhibitory concentration; SEM, standard error of the mean, N/A, no activity observed. Data are representative of two independent experiments.

Interestingly, replacement of Lys^19^ by Ala had the greatest impact on activity, reducing the antimicrobial activity by 128-fold: 667 ng/μL was required to inhibit growth of the indicator strain as no turbidity was observed in the overnight culture, compared to 5.2 ng/μL for the wild type. The control we used was the untransformed *L. plantarum* WCFS1 with 10 mM ammonium acetate, which demonstrated no MIC. This control confirms that the inhibitory activity was due to the respective bacteriocins alone. These MIC results identify an important functional role for Phe^8^, Trp^45^ and Lys^19^ three highly conserved residues identified in the sequence analysis of circular bacteriocins (Figure 4). The MIC results are in agreement with previous findings that conserved aromatic and cationic residues are important for antimicrobial activity ^10^ of plantacyclin B21AG and by extension, other circular bacteriocins which share these features.

The observed masses of the wild type plantacyclin B21AG and its variants are listed in Table 4. Notably, a mass was not identified for the plantacyclin B21AG Lys^19^Ala mutant, despite multiple attempts using fresh cultures and purifications. This mutant had a reduced antimicrobial activity compared with wildtype (128-fold reduction). We suggest that substitution of Lys^19^ with Ala may result in incorrect cellular processing or secretion, instability and/or reduced stability of the mutant plantacyclin B21AG. A possible explanation is that the N-C terminal circularisation may be compromised (this circularisation is a key feature of stability ^30,39^).

**Table 4.** MALDI-TOF mass determination of plantacyclin B21AG-WT and mutants. Also shown is the expected size and mass differences.

| *L. plantarum* WCFS1 Sample | Expected molecular weight if circularised | MALDI-TOF results | MALDI-TOF mass minus expected mass |
| --- | --- | --- | --- |
| Plantacyclin B21AG-WT | 5667.7 | 5672.1 | 4.4 |
| Plantacyclin B21AG-Phe8Ala | 5591.6 | 5590.4 | -1.2 |
| Plantacyclin B21AG -Lys19Ala | 5610.6 | Not identified | N/A |
| Plantacyclin B21AG -Trp45Ala | 5552.6 | 5550.9 | -1.7 |
| Untransformed (-) | N/A | Not identified | N/A |
| 10 mM ammonium acetate (-) | N/A | Not identified | N/A |

## Conclusions

We report the crystal structure of a circular bacteriocin from the food grade *Lactobacillus plantarum* B21. Structural comparison of plantacyclin B21AG and other circular bacteriocins confirms that these antimicrobial peptides share a highly conserved core fold and secondary structure composition, despite low sequence identity. In the structures solved in aqueous conditions, the N and C ligation point is buried in the hydrophobic core of the globular structure and positioned within an α helix. Sequence alignment of plantacyclin B21AG with other bacteriocins that have been characterised to be circular revealed highly conserved motifs including polycationic regions predicted to be surface exposed that could be involved in membrane interaction, binding and destabilisation. Proximally located anionic residues could be involved in pore selectivity that drives cation flow out of the target cell. Another common motif was the presence of conserved aromatic residues near the N and C terminal ends of helices 4 – 5. These could be involved in membrane solubilisation, penetration and channel formation. Through multiple lines of evidence including sequence analysis, structure analysis, site-directed mutagenesis and phenotypic activity, we demonstrated the importance of these evolutionarily conserved residues. In one case, removal of Lys^19^ resulted in 128-fold reduction of antimicrobial activity compared to the wild type plantacyclin B21AG. A problem encountered was the inefficient transformation efficiency of *L. plantarum* WCFS1 when transforming pTRKH2 backbone plasmids. Future work could focus on improving the efficiency by utilising different plasmid backbones and optimising the transformation protocol, to enable more rapid testing of other variants. Despite this limitation, we demonstrated that three aromatic and cationic residues have key functional roles in the antimicrobial activity of plantacyclin B21AG. Sequence and structure analysis suggest that several other conserved residues likely have functional roles. These include Arg^6^, His^11^ as well as the C and N terminal residues, and could be tested in a similar manner when the transformation efficiency is improved.

## Methods

### Expression and purification of plantacyclin B21AG

*Lactobacillus plantarum* B21 was inoculated in 1 L of de Man, Rogosa & Sharpe (MRS) broth using a 2% inoculum (v/v). The culture was incubated for 16 – 18 hrs at 30°C without shaking. The bacteriocin plantacyclin B21AG was purified from the culture supernatant through a four-step protocol, i.e. concentration of cell free supernatant, extraction with water-saturated butanol, desalting through PD10 and cation exchange Fast Protein Liquid Chromatography (FPLC). After overnight incubation, the culture was pelleted at 10,000 x g for 10 min. The resulting cell free supernatant (CFS) was concentrated through a 10 kDa polyethersulfone membrane disc using an Amicon® Stirred Cell (Merck Millipore, Germany) to a final volume of 50 mL. The concentrated CFS was then extracted twice with ½ volume of water saturated butanol. The butanol fraction containing plantacyclin B21AG was dried under a fine stream of nitrogen to remove the butanol. The dried butanol fraction was then redissolved in 20mM of sodium phosphate buffer (pH 6) and desalted using a NAP10 desalting column prepacked with Sephadex G-25 resin (GE healthcare, USA). Finally, the desalted protein fraction was purified using the Uno S6 prepacked monolith cation exchange column (12 × 53 mm, 6 mL, Bio-Rad, USA). The eluted fraction containing purified bacteriocin was concentrated to 4.6 mg/mL in an Amicon centrifugal filter concentrator with a 3 kDa cutoff membrane (Millipore) in buffer comprising 20 mM sodium phosphate (pH 6) for subsequent protein crystallisation.

### Protein crystallisation and data collection

Protein crystallisation was performed at the UQ ROCX crystallisation facility at the Institute for Molecular Bioscience, University of Queensland. JCSG-*plus*™ HT-96 commercial screens (Molecular Dimensions, USA) was used to screen for crystallisation conditions using the hanging drop vapour diffusion method. Briefly, plantacyclin B21AG was dissolved in water at a concentration of 2.3 mg/mL. Each hanging drop comprised 100 nL of purified bacteriocin and 100 nL of crystallisation solution and were set up using a Mosquito crystallisation robot (TTP Labtech, UK) in a 96-well format. The crystallisation trays were incubated at 20°C in (and imaged using) a RockImager 1000 (Formulatrix, USA).

After 12 hours incubation, a single crystal formed in condition F10 of JCSG screen containing 1.1 M sodium malonate, 0.1 M HEPES buffer pH 7.0 and 0.5% v/v Jeffamine® ED-2003. A single crystal was harvested after 13 days of incubation. X-ray diffraction experiments were performed at the micro-focus beamline (MX2) of the Australian Synchrotron ^64^. Crystals were flash-cooled and stored in liquid nitrogen before transferring to a stream of nitrogen gas at 100 K at the beamline. X-ray diffraction data were collected at a wavelength of 0.9537 Å using an EIGER-16 M detector at distance of 230.028 mm and with 0.1° oscillation and 0.1 s exposure of 0% attenuated beam per frame. 3600 frames of the data set were collected in 36 sec. The data were processed and scaled using XDS^65^ and Aimless ^66^, respectively.

### Crystal structure determination and refinement

Attempts to solve the X-ray crystal structure of plantacyclin B21AG by molecular replacement (MR) using the NMR structure of acidocin B did not yield a solution. Instead, the phases for plantacyclin B21AG crystal structure were solved by molecular replacement (MR) method with a short model helix structure of 8 amino acids (polyalanine) as a search model, using the program PHASER in the CCP4 suite ^67,68^. The program placed six helices in an asymmetric unit with LLG of 393. The six helical model was used in SHELXE^69-71^ for density modification and polyala model building. The polyala model (109 residues in two chain) resulting from SHELXE was provided to MR phasing protocol of Auto-Rickshaw^72,73^ server (http://www.embl-hamburg.de/Auto-Rickshaw) for phase improvement, model building and sequence docking. Within the software pipeline, MR was skipped and the partial model was refined in CNS ^74^ for B-factor and positional refinement. Further refinement in REFMAC5 ^75,76^ resulted in R/Rfree of 38.7%/40.1%. Density modification was performed in PIRATE ^77^ and the resulting phases were used in ARP/wARP^78^ for automated model building. The above density modification and model building procedure were repeated twice, which resulted in a model containing 112 residues with docked sequence in two chains. The last round of refinement (using REFMAC5) gave R/Rfree of 25.7%/30.5%. At this stage, the model was used for manual rebuilding and refinement. 56 water molecules were included where the mFo-nFc difference electron density showed a peak above 3σ and the modelled water made stereochemically reasonable hydrogen bonds. Final refinement was performed with REFMAC5. A Ramachandran plot showed that 99.1% of the residues were in the preferred regions and 0.9% were in the allowed regions. The quality and geometry of the final structure were evaluated and validated using wwPDB validation system ^79^. Figures were prepared using PYMOL ^55^. Data collection and refinement statistics are presented in Table 1. The final refined model has been deposited in the Protein Data Bank with the code 6WI6 using wwPDB OneDep system ^80^.

### Structural analysis

Superposition of Chain A and Chain B of Plantacyclin B21AG was performed using WinCoot 0.8.9.2 ^81^. The percentage of solvent exposure of each amino acid residue was calculated using GETAREA 1.0 beta (http://curie.utmb.edu/getarea.html) ^44^, a web service provided by the Sealy Center for Structural Biology at the University of Texas Medical Branch. The electrostatic potential surface map was calculated using the APBS functionality ^82^ in the PDB2PQR web server (version 2.1.1) (http://nbcr-222.ucsd.edu/pdb2pqr_2.1.1/) ^83^ and the figures generated using Chimera 1.14 ^84^. The isoelectric focusing point of the proteins was computed using the Compute pI/MW tool in ExPASy (https://web.expasy.org/compute_pi/). Structural alignment between circular bacteriocins was performed using the Align function in PYMOL ^55^.

### Circular bacteriocin sequence analysis

Plantacyclin B21AG sequence was aligned against sequences of bacteriocins previously characterised to be circular. The sequences were mined from NCBI database and aligned using Clustal Omega (https://www.ebi.ac.uk/Tools/msa/clustalo/) ^85^. The alignment was manually edited to remove gaps for figure clarity and exported to fasta format, which was converted to phylip format using Sequence Conversion (http://sequenceconversion.bugaco.com/converter/biology/sequences/fasta_to_phylip.php). This was used as input into RAxML (raxmlHPC-PTHREADS-SSE3 version 8.2.10) ^86^ using the following parameters for ML + rapid bootstrap analysis with 100 replicates: -T 2 -f a -x 285 -m PROTGAMMABLOSUM62 -p 639 -N 100 The bipartitions output file was used in FigTree version 1.4.4 (http://tree.bio.ed.ac.uk/software/figtree/) for viewing/manipulation.

Jpred 4 ^87^ (http://www.compbio.dundee.ac.uk/jpred/, date accessed: 2/9/18) was used for prediction of secondary structures of sequences without confirmed structures. The Jnet prediction algorithm was initially run, then to determine the secondary structures around the C and N termini, protein sequences were rearranged, and run again.

### Site-directed mutagenesis

Plasmid construct pCycB21^42^ was used as a basis for mutagenesis studies. 998 bp inserts containing site-directed mutations were generated via DNA synthesis (Biomatik), flanked with *BamH*I/*Pci*I restriction sites and 6 base pair (bp) protection bases. These were subcloned into pCycB21, replacing the wild type (WT) sequence. The details of each construct are found in Table 5.

**Table 5.**
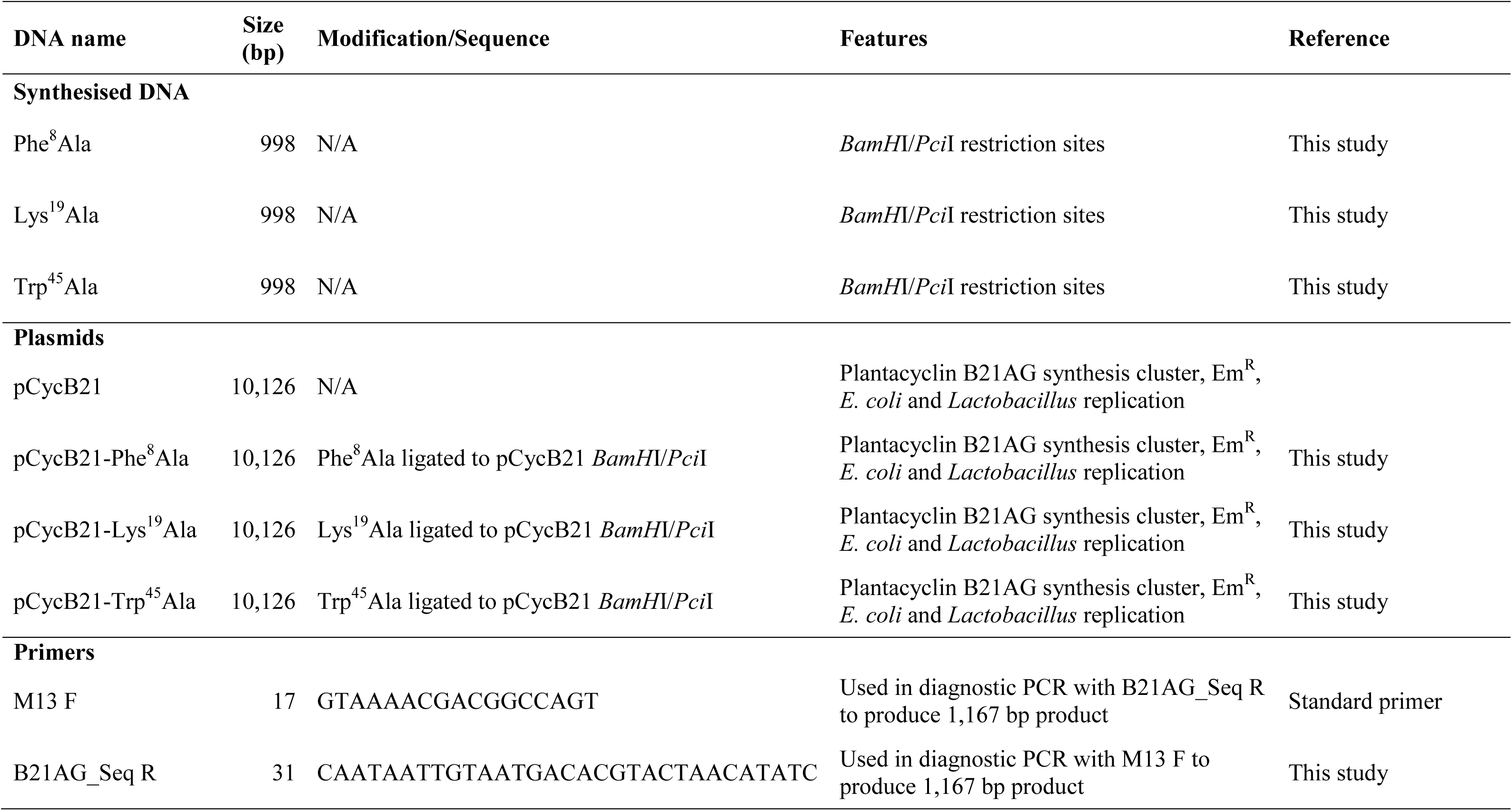
DNA and plasmids used in this study

### Transformation into plantacyclin B21AG-deficient strain

As previously shown, *Lactobacillus plantarum* WCFS1 shares 99% genome identity to original plantacyclin B21AG producer *L. plantarum* B21 ^88^. The generated constructs were transformed into *Lactobacillus plantarum* WCFS1 (ATCC BAA-793) ^42^ via electroporation ^89^ with some alterations. Briefly, 0.1 mL of an overnight culture was inoculated into 5 mL MRS media (OXOID) containing 1% glycine and 0.75 M sorbitol. Cells were grown at 37 °C until mid-exponential phase (OD_600_ of 1.5 - 3) then centrifuged at 4000 x g for 5 minutes at 4°C. From this point on, cells were kept on ice and centrifuged as above. Cells were then washed twice with a 10:1 volume of transformation solution containing 950 mM sucrose and 3.5 mM MgCl_2_. Cells were resuspended in 80 μL of transformation buffer and 500 ng of plasmid DNA was added to cells. This mixture was then added to 0.2 cm cuvette (Bio-Rad) and electroporated with a Gene Pulser (Bio-Rad) with the following conditions: 2 kV, 25 μF and 400 Ω. If the time constant was ≥10, another 500 ng of DNA was added, and the electrical pulse repeated. 1 mL of MRS containing 100 mM MgCl_2_ and 0.5M sucrose was used to quickly rescue cells from cuvette. The mixture was added to 1.5 mL microtubes and cells were incubated at 37°C for 3 hours, then 200 μL were spread onto MRS agar plates containing 15 μg/mL of erythromycin (Sigma-Aldrich). Plates were grown at 37 °C for 48 hours and colonies screened via erythromycin resistance phenotype and Polymerase Chain Reaction (PCR). Sanger sequencing was performed on the PCR products using both primers to confirm the transformants, at the Griffith University DNA Sequencing Facility, QLD, Australia.

### PCR

PCR was performed using GoTaq Green Master Mix (Promega) as per manufacturer’s instructions. Thermocycling conditions were as follows: Initial 2-minute melting step of 95°C, followed by 35 cycles of: 95 °C for 15 seconds, 52 °C for 15 seconds, 72 °C for 1 minute 20 seconds. A final extension of 72°C for 5 minutes was performed.

### Partial-purification of *L. plantarum* WCFS1-derived bacteriocins

After *L. plantarum* WCFS1 transformants had been confirmed, they were grown for 24 hours in 100 mL MRS cultures containing 15 μg/mL erythromycin. Untransformed *L. plantarum* WCFS1 cells were also grown overnight in MRS without antibiotics to serve as a negative control. Cells were centrifuged at 8,000 x g for 10 minutes, then supernatant concentrated to 2 mL using 3 kDa Amicon Ultra-15 Centrifugal Filter Units (Merck). Supernatant was then washed 3 times with 15 mL 10 mM ammonium acetate to remove residual erythromycin. Concentrated supernatant was then n-butanol extracted as above and then removed in a rotary evaporator. The dried fraction was then resuspended in 10 mM ammonium acetate. Concentration of the bacteriocin from the partial purification was calculated using the extinction coefficient ^90^ and 280 nm absorbance value using the ProtParam calculator http://www.protparam.net/index.html. Concentration was also evaluated with a Bradford assay, using bovine serum albumin as a standard to increase confidence in values generated (data not shown).

### Mass analysis using MALDI-TOF MS

Samples were analysed in a buffer containing 10 mM ammonium acetate with 0.1% formic acid. Analysis was performed on a ABSciex MALDI 5800 at the Mass Spectrometry Facility, University of Queensland, Centre for Clinical Research, QLD, Australia. A Bacterial Test Standard (Bruker) was used to calibrate the instrument. Partially purified supernatant from *L. plantarum* WCFS1 untransformed (-) was used as a negative control along with 10 mM ammonium acetate with 0.1% formic acid.

### Minimum inhibitory concentration assays of partially purified site-directed mutants

After partial-purification, the minimum inhibitory concentration (MIC) of the plantacyclin B21AG-WT and mutants were evaluated. WT refers to the bacteriocin produced by *L. plantarum* WCFS1 transformed with pCycB21 rather than native producer *L. plantarum* B21, herein this manuscript referred to as plantacyclin B21AG.

MIC assays were performed in clear 96 well plates (Greiner). 150 μL MRS containing 3 × 10^5^ CFU/mL of indicator strain *L. plantarum* A6 ^91,92^ were added to wells. Partially purified bacteriocin samples were examined by making dilutions in 150 μL 10 mM ammonium acetate, then added to the wells. 1:2 serial dilutions of the bacteriocin were performed, evaluating each plantacyclin B21AG mutant from 1000 ng/μL to 5.2 ng/μL. Supernatant from *L. plantarum* WCFS1 untransformed (-) and 10 mM ammonium acetate without bacteriocin were used as a bacteriocin negative controls. 10 mM ammonium acetate without bacteriocin or indicator strain was used as a blanking control. Plates were incubated at 37 °C for 24 hours. Assays were performed in biological duplicate. Plates were read using a Synergy 2 (BioTek) plate reader at 600 nm absorbance. MIC value was determined at the minimum concentration of bacteriocin where no growth of *L. plantarum* A6 occurred.

## Data availability

The plantacyclin B21AG crystal structure coordinates and structure factors are available in the Protein Data Bank repository with a PDB code 6WI6, [https://www.rcsb.org/].

## Acknowledgements

This study was supported by Griffith University Postdoctoral Fellowships. We thank Karl Byriel, the manager of the University of Queensland Remote Operation Crystallization and X-ray Diffraction Facility (UQROCX) that we accessed for protein crystallization, crystal imaging and X-ray diffraction. We thank the University of Queensland Institute for Molecular Bioscience’s (IMB) Mass Spectrometry Facility for access to MALDI-TOF-MS. This research was undertaken in part using the MX2 beamline at the Australian Synchrotron, ANSTO and made use of the Australian Cancer Research Foundation (ACRF) detector and we are grateful for access to the Synchrotron, for data collection and processing.

## Author contributions

A.T.S, J.L.M., M.C.G., B.R. and B.V conceived and designed the experiment(s); M.C.G., B.V., R.M.M. and G.K. conducted the experiment(s), S.P., M.C.G. and B.V. analysed the data. J.L.M. provided access to instrumentation at Griffith University. M.C.G and B.V. wrote the main manuscript text. All authors reviewed the manuscript.

